# Root-exuded secondary metabolites can alleviate negative plant-soil feedbacks

**DOI:** 10.1101/2023.04.09.536155

**Authors:** Valentin Gfeller, Lisa Thönen, Matthias Erb

## Abstract

Plants can suppress the growth of other plants by modifying soil properties. These negative plant-soil feedbacks are often species-specific, suggesting that some plants possess resistance strategies. However, the underlying mechanisms remain largely unknown. Here, we investigated if and how benzoxazinoids, a class of dominant secondary metabolites that are exuded into the soil by maize and other cereals, help plants to resist negative plant-soil feedbacks. We find that three out of five tested crop species suppress maize performance relative to the mean across species. This effect is partially alleviated by the plant’s capacity to produce benzoxazinoids. Soil complementation of benzoxazinoid-deficient mutants with purified benzoxazinoids is sufficient to restore the protective effect. Sterilization and reinoculation experiments suggest that benzoxazinoid-mediated protection acts via changes in soil microbes. Substantial variation of the protective effect between experiments and soil types illustrates that the magnitude of the protective effect of benzoxazinoids against negative plant-soil feedbacks is context dependent. In summary, our study demonstrates that plant secondary metabolites can confer resistance to negative plant-soil feedbacks. These findings expand the functional repertoire of plant secondary metabolites, reveal a mechanism by which plants can resist suppressive soils, and may represent a promising avenue to stabilize plant performance in crop rotations in the future.

## Introduction

Plants constantly interact with their soil environment. They change the biotic and abiotic soil attributes, which then, in turn, alters the performance of proceeding plants. These so-called plant-soil feedbacks can either enhance or lower the performance of the following plant (Bever *et al*., 1997; van der Putten *et al*., 2013). Plant-soil feedbacks are involved in many ecological processes including vegetation succession, plant invasion, and maintenance of species diversity (van der Putten *et al*., 1993; Klironomos, 2002; van der Putten *et al*., 2013; Teste *et al*., 2017). In agriculture, they have been used for centuries to mitigate negative impacts of monocropping. Efforts to translate ecological knowledge from plant-soil feedback research into improved crop rotations are now needed (Mariotte *et al*., 2018). Such evidence based crop rotation design represents a promising avenue towards more sustainable agriculture (Dias *et al*., 2015; Mariotte *et al*., 2018).

Crop rotations are best studied for their long-term benefits. Over years of cultivation, crop rotations are capable of increasing soil health and suppressing weeds, pathogens, pests, and insects (Brust & King, 1994; Karlen *et al*., 1994; Chen *et al*., 2001; McDaniel *et al*., 2014; Tiemann *et al*., 2015; Leandro *et al*., 2018). In addition, making cropping systems more diverse also makes them more resilient against adverse growth conditions and weather extremes (Bowles *et al*., 2020), this will be of importance to alleviate adverse impacts of global change. Soil conditioning by a given crop species can alter the growth, defence, yield, and soil processes of the following crop plant (Sieling & Christen, 2015; McDaniel *et al*., 2016; Benitez *et al*., 2017). Benitez and colleagues, for example, showed that precrop identity alters the microbial communities in the rhizosphere of maize seedlings and affects their performance. Given that plant-associated microbes are known to be important determinants for plant health (Berendsen *et al*., 2012), it is tempting to hypothesize that changes in maize seedling performance are driven by precrop-dependent microbiomes.

Negative plant-soil feedbacks can be triggered by reduced nutrient availability, accumulation of soil-borne pathogens, depletion of beneficial microbes and changes in soil chemistry (Bennett & Klironomos, 2019; Schandry & Becker, 2020). The relative contribution of these mechanisms depends on the environmental context (Smith-Ramesh & Reynolds, 2017). Furthermore, the direction and magnitude of plant-soil feedbacks is also species- and variety-specific (Bever, 1994; Cadot *et al*., 2021a; Awodele & Bennett, 2022), indicating that distinct resistance strategies exist. How plants resist negative plant-soil feedbacks is, however, largely unknown. Understanding this process may help to increase the stability of crop rotations.

Root exudates shape the rhizosphere microbiome (Sasse *et al*., 2018). For benzoxazinoids and flavones these changes were linked to the performance of succeeding conspecific plants (Hu *et al*., 2018b; Yu *et al*., 2021). Through plant-soil feedbacks, the trait of benzoxazinoid exudation in maize also affects wheat performance under controlled conditions and improves wheat yield under field conditions (Cadot *et al*., 2021a; Gfeller *et al*., 2022). Considering that benzoxazinoids can structure the rhizosphere microbiome of maize already at the seedling stage (Cotton *et al*., 2019; Kudjordjie *et al*., 2019), that benzoxazinoids can act fungistatic against soil pathogens (Wilkes *et al*., 1999; Martyniuk *et al*., 2006), and that benzoxazinoids can attract a *Pseudomonas putida* strain that potentially induces plant resistance (Neal *et al*., 2012; Neal & Ton, 2013), we hypothesize that they could also increase crop rotation stability by alleviating negative plant-soil feedbacks.

Benzoxazinoids, a class of indole-derived plant secondary metabolites, are well known for their multifaceted bioactivities (Niemeyer, 2009). They are most prevalent in grasses, including agronomically important crops such as maize, wheat, and rye (Frey *et al*., 2009). Besides their effects on microbes, they are well known as defence metabolites against insects and pathogens (Niemeyer, 2009), and they can act as signalling molecules (Ahmad *et al*., 2011). Further, it is known that benzoxazinoids have chelating properties leading to improved iron acquisition (Hu *et al*., 2018a).

Here, we investigate the role of benzoxazinoids in tolerating crop rotation legacies. We hypothesize that benzoxazinoid exudation into the rhizosphere reduces negative plant-soil feedbacks caused by certain precrops. By growing wild-type and benzoxazinoid-deficient *bx1* mutant maize, we examined how benzoxazinoids alter soil legacy effects of different precrops. In several plant-soil feedback experiments, we tested if the direction and/or magnitude of these feedbacks change with different precrops, soils, and/or response maize lines. Through chemical complementation and sterilization experiments, we further assessed the underlying mechanism. We found that benzoxazinoids increase crop rotation stability through root exudation and soil biota dependent mechanisms, while also unmeasured factors contributed to the outcome of some experiments.

## Materials and Methods

### Plant material

To investigate the effect of maize benzoxazinoids in resistance to negative plant-soil feedbacks, we selected five plant species as precrops and two maize lines with their corresponding benzoxazinoid deficient mutant as response plants. We selected a genetically diverse set of precrops belonging to four different families, all of them commonly cultivated in crop rotations with maize: *Glycine max* cv. green shell (soybean), *Medicago sativa* (alfalfa), *Brassica napus* (rapeseed), *Phacelia tanacetifolia* (lacy phacelia), and *Triticum aestivum* cv. Claro (winter wheat). *G. max*, *M. sativa*, and *P. tanacetifolia* seeds were obtained from Sativa Rheinau AG (Switzerland), *B. napus* seeds were purchased online (www.saemereien.ch), and *T. aestivum* seeds were kindly provided by Saatzucht Düdingen (Switzerland). To ensure nodulation, *G. max* seeds were inoculated with rhizobia (LegumeFix, Sativa Rheinau AG) according to the supplier’s recommendations. The maize lines W22 and B73 were selected as response plants, since for them benzoxazinoid-deficient *bx1* mutants are available (Tzin *et al*., 2015; Maag *et al*., 2016). We did not include bare soil as a treatment, but instead used the mean performance of maize across all precrops as an ecologically meaningful baseline to assess whether individual precrops cause significant positive or negative plant-soil feedbacks.

### Soil material

Feedback experiments were conducted in field soil (clay loam) collected in three batches at the Agroscope field station in Changins (Switzerland). For the initial precrop screening, soil was sampled on field parcel 29. For all the other experiments, soil was sampled in two batches on another filed, parcel 30. An additional soil (silt loam), referred to as Q-Matte, was collected from a grassland site near Bern (Switzerland) and was used to test for soil-specific effects. Collected soil was sieved (10 mm mesh size), completely homogenized, and stored at 4 °C before utilization. Soils were characterized in previous publications (Hu *et al*., 2018b; Cadot *et al*., 2021a).

### Plant growth

Experiments were performed in walk-in climate chambers under controlled conditions (day length: 14 h; temperature: 22 °C/18 °C; humidity: 60 %; light: ∼ 550 µmol m^-2^s^-1^). In the conditioning phase the precrops were grown in 2 L pots (Rosentopf Soparco 2.0 l; Hortima, Switzerland) for six weeks, followed by the maize feedback phase in either 2 L or 1 L pots (Rosentopf Soparco 1.0 L; Hortima, Switzerland) for six weeks or four weeks, respectively. To avoid the roots from growing out of the pot, fleece (Geotex; Windhager, Austria) was placed at the bottom of each pot, before filling with soil. Pots were subsequently put in the climate chamber to acclimatize for at least one day before sowing. For each precrop an excess of seeds was sown and thinned out to two plants per plot after one week, except for the fast-growing soybean, where we only kept one plant. Plants were watered as needed, and once a week, 100 mL of a nutrient solution (0.2% [w/v]; Plantaaktiv Typ K, Hauert, Grossaffoltern, Switzerland) supplemented with iron (1 ‰ [w/v]; Sequestrene rapid, Maag) was supplied in the conditioning phase and increased to 0.02 % [w/v] Sequestrene in the feedback phase (unless otherwise stated). Pots were randomly arranged in the climate chamber and re-randomized on a weekly basis. At precrop harvest, shoot biomass was collected and dried until constant weight at 80 °C, before dry weight was determined on a microbalance. After removing the root system, the remaining soil was sieved (10 mm mesh size), homogenized within each precrop (in all but one experiment), and used for the feedback phase. In the feedback phase, pot preparation was performed identical to the conditioning phase and maize seeds were sown. At harvest, plant height was measured and, in some experiments, chlorophyll content was determined by averaging nine measurements equally distributed along the youngest fully opened leaf by means of a SPADE-502 chlorophyll meter (Konica Minolta, Japan), and maize biomass was weighted after drying at 80 °C until constant weight. See **Fig. S1** for specific information on pot size, length of feedback period, and soil treatments between conditioning and feedback phase.

### Screening benzoxazinoid-mediated resistance to plant-soil feedbacks

To examine the role of benzoxazinoids in plant-soil feedback responses to different precrops, wild-type B73 maize and *bx1* mutants in the B73 background were grown for six weeks in 2 L pots. To analyse benzoxazinoid exudation, we sampled the soils of three wild-type pots and one *bx1* pot per precrop species at the end of the experiment. We sieved the soil through a 10 mm sieve, again sieved a subset of this soil through a test sieve (5 mm, Retsch, Haan, Germany), and filled 25 mL soil into a 50 mL centrifuge tube. The tubes were then stored at – 80 °C until further processing (see below).

### Examination of soil type, maize line, and plant age dependency

To test if benzoxazinoid-mediated resistance to negative plant-soil feedbacks depends on the soil type, maize line, and/or plant age, we performed a feedback experiment comparing B73 and W22 maize in soil from Changins. For the W22 genetic background, we also compared two soils, Changins and Q-Matte. Based on their strong positive and negative effects on maize growth, we focused on *M. sativa* and *T. aestivum* as representative precrops for these experiments. In the response phase, maize was grown for six weeks in 2 L pots. The following adjustments were made compared to all other experiments: (i) the experimental units were kept separate from conditioning to feedback phase without mixing, and (ii) fertilization was maintained low during precrop conditioning and maize response. 100 mL of a nutrient solution (0.2% [w/v]; Plantaaktiv Typ K, Hauert, Grossaffoltern, Switzerland) supplemented with iron (1 ‰ [w/v]; Sequestrene rapid, Maag) was supplied on a weekly basis.

### Mechanistic experiments

To evaluate the mechanism by which benzoxazinoids protect maize plants against negative plant-soil feedbacks, we performed a series of complementation, sterilization, and re-inoculation experiments. To test for robustness, we performed these experiments three times. Experiment 1 and 2 were performed in the same soil batch as the previous experiment, whereas for experiment 3, we collected fresh soil from the same location (Changins, **Fig. S1**), as the initial soil batch was depleted. In all three experiments, feedbacks were conducted with W22 maize in 1 L pots for four weeks.

### Soil complementation with benzoxazinoids

To test if the observed differences in performance of wild-type and *bx1* mutant plants are triggered by benzoxazinoids exuded into soil matrix, we externally applied a mixture of benzoxazinoids to *bx1* mutant plants growing in *T. aestivum* conditioned soil. In the three experiments, benzoxazinoids were applied in three different concentrations (**Fig. S5a,b**). Benzoxazinoid levels in the soil were determined at the end of each experiment, to estimate the effectiveness of our treatment. AMPO, a stable degradation product of benzoxazinoids, was increased in the soil following benzoxazinoid complementation. All other compounds were only detected in trace amounts in complemented soils, suggesting rapid degradation in the absence of a constant emitter (**Fig. S5c-e**). Compared to levels in the soil of wild-type plants, AMPO concentrations in complemented *bx1* mutant soil were very low in the first experiment, where we applied 50 ug of benzoxazinoids per week and pot. We thus increased our complementation dose in the second experiment to 5.5 mg, resulting in AMPO concentrations that were higher than in soils of wild-type plants. For the third experiment, we thus used an intermediate dose, 1.6 mg, resulting in wild-type levels (**Fig. S5c-e**). For complementation, benzoxazinoids were purified form four day old seedlings (see below), dissolved in deionized water, and 5 mL of this solution was pipetted to the *bx1* plants every three days, starting two days after sowing (at germination). Control *bx1* plants and wild-type plants were supplied with the same amount of deionized water. To investigate benzoxazinoid accumulation in the pots, soil was sampled as described above for a random subset of plants.

### Sterilization and reinoculation experiments

To evaluate if soil biota are driving the positive effects of benzoxazinoids on plant growth, *T. aestivum* conditioned soils were X-ray sterilized (20-60 kGy; Steris, Däniken, Switzerland). In the feedback phase wild-type and *bx1* mutant plants were grown in unsterilized, sterilized, and re-inoculated soil. Re-inoculation was achieved by complementing 95 % of sterilized soil with 5 % of unsterilized (living) *T. aestivum* conditioned soil and homogenizing thoroughly. All soils were acclimatized for one week in the climate chamber before sowing. All plants were watered with autoclaved tab water. To further investigate the relative contribution of the soil biota and abiotic soil attributes, we also tested for precrop-specific inoculation effects. Therefore, we included four additional soil conditions consisting of unsterilized *M. sativa* soil, sterilized *M. sativa* soil, and sterilized *M. sativa* soil inoculated with either unsterilized *M. sativa* or *T. aestivum* soil.

### Purification of benzoxazinoids for complementation

To purify benzoxazinoids, 40 g of maize seeds (var. Akku) were placed in a 1 L glass beaker and soaked in autoclaved water for 14 h. Kernels were washed twice a day and harvested after four days. Soaking and growth took place in the dark at 26 °C. During harvest, kernels were immediately put into a blender (MioStar Beld 600s; Migros, Switzerland) prefilled with 600 mL methanol (MeOH), blended at maximum speed for five minutes, and passed through a filter paper (Grade 1; Whatman, GE Healthcare Live Sciences, USA). Next, we removed MeOH and water in the extracts by evaporation (40 °C; rotary evaporator), followed by freeze drying. The dry material was dissolved in MeOH, bound on silica (0.062-0.2 mm), evaporated to dryness, and compounds were separated on a flash chromatography purification system (CombiFlash RF+, Teledyne ISCO, USA) in two subsequent runs, where benzoxazinoids were detected at wavelength 254 nm. The first run was performed on a 120 g RediSep Silica column at a flow rate of 85 ml/min, with chloroform (stab./EtOH; solvent A) and MeOH (solvent B) as solvents. The elution profile was as follows: 0-2 min, 0-13 % B; 2-6 min, 13-16 % B; 6-7.5 min, 16% B; 7.5-9.6 min, 16-33.6%; 9.6-12.7 min, 33.6-58% B, and kept at 58 % B. The second run was performed on a 40 g RediSep Silica column at a flow rate of 40 ml/min with the same solvents and the following elution profile: 0-2 min, 0-15 % B; 2-3 min, 15 % B; 3-8.7 min, 25-30 % B, and kept at 30% B. The fractions containing benzoxazinoids were evaporated on a rotary evaporator (40 °C), sterile filtered through a PTFE 0.20 (ChromafilXtra; MN, Germany) filter, and evaporated to dryness. To crystallize the benzoxazinoid mixture, the compounds were dissolved in deionized water and lyophilized. The resulting white powder was used for complementation and an aliquot was characterized on a UHPLC-MS system (see below).

### Analysis of benzoxazinoids

To analyse benzoxazinoids and break down products, soil was sampled as described above. The frozen 50 mL centrifuge tubes containing the soil were thawed before the soil was dissolved in 25 mL acidified MeOH/H_2_O (70:30 v/v; 0.1% formic acid). The suspension was placed on a rotary shaker for 30 minutes at room temperature, followed by sedimentation of the soil by centrifugation (5 min, 2000 g). The supernatant was filtered (Filter paper, Grade 1; Size: 185 mm; Whatman, GE Healthcare Live Sciences), a 1 mL aliquot of the filtrate was transferred into a 1.5 mL centrifuge tube, centrifuged (10 min, 19000 g, 4 °C), and the supernatant was sterile filtered (Target2TM, Regenerated Cellulose Syringe Filters. Pore size: 0.45 µm; Thermo Scientific) into a glass tube for analysis.

Benzoxazinoids extracted from soils and the purified benzoxazinoid mixture from germinated maize kernels were analysis as described before (Gfeller *et al*., 2022). In short, an Acquity UHPLC system coupled to a G2-XS QTOF mass spectrometer equipped with an electrospray source and piloted by the software MassLynx 4.1 (Waters AG, Baden-Dättwil, Switzerland) was used. Absolute quantities were determined through standard curves of pure compounds. For that, MBOA (6-methoxy-benzoxazolin-2(3H)-one) was purchased from Sigma-Aldrich Chemie GmbH (Buchs, Switzerland). DIMBOA-Glc (2-O-β-D-glucopyranosyl-2,4-dihydroxy-7-methoxy-2H-1,4-benzoxazin-3(4H)-one) and HDMBOA-Glc (2-O-β-D-glucopyranosyl-2-hydroxy-4,7-dimethoxy-2H-1,4-benzoxazin-3(4H)-one) were isolated from maize plants in our laboratory. DIMBOA (2,4-dihydroxy-7-methoxy-2H-1,4-benzoxazin-3(4H)-one), HMBOA (2-hydroxy-7-methoxy-2H-1,4-benzoxazin-3(4H)-one), and AMPO (9-methoxy-2-amino-3H-phenoxazin-3-one) were synthesized in our laboratory.

### Statistical analysis

All statistical analysis were conducted in R version 4.1.2. (R Core Team, 2021). Data management and visualisation was facilitated with the *tidyverse* package collection (Wickham *et al*., 2019). Phenotypic data was analysed by analysis of variance (ANOVA) unless otherwise stated. For that, statistical assumptions such as normal distribution and homoscedasticity of error variance were visually checked. If treatments showed unequal variance, a generalized least squares model was fitted using the gls() function of the *nlme* package (Pinheiro *et al*., 2021). Differences in estimated marginal means (EMMs) were analysed by pairwise comparison with the emmeans() function of the *emmeans* package and false discovery (FDR) corrected *p* values were reported (Benjamini & Hochberg, 1995; Lenth, 2022). To evaluate the effect of individual precrop soil conditioning on the subsequent maize performance, one-sample t-tests were performed for every precrop, comparing its biomass production to the overall mean of all precrops. To test for differences in benzoxazinoid production between wild-type and *bx1* mutant maize as well as validation of complementation success, Wilcoxon rank-sum tests were performed. Differences in weight gain between the precrops *T. aestivum* and *M. sativa* were tested with Welch’s two-sample *t*-test. To test at what point in time the growth increase of wild-type plants relative to *bx1* mutant plants became statistically significant, Welch’s two-sample *t*-tests were performed and FDR corrected *p* values were reported. The endpoint analysis of the time series experiment was also analysed by Welch’s two-sample *t*-tests.

## Results

### Benzoxazinoids enhance resistance to negative plant-soil feedbacks

To test whether benzoxazinoids enable maize plants to cope with negative plant-soil feedbacks, we grew five crop species for six weeks under controlled conditions, followed by a feedback phase with wild-type (B73) and benzoxazinoid-deficient *bx1* mutant maize. Biomass accumulation after six weeks ranged from 3.4 g to 10.5 g dry weight (**Fig. S2a**). After harvesting the conditioning plants, conditioned soils were sieved, and wild-type and *bx1* mutant maize was sown (**Fig. S1**). After six weeks of growth, we observed differences in biomass accumulation of maize depending on the precrop (**Fig. 1a**). Maize plants accumulated significantly less biomass on soils conditioned by *T. aestivum, P. tanacetifolia,* and *B. napus* compared to the overall mean of all precrops (*p* < 0.001), indicating a negative plant-soil feedback. In contrast, maize plants accumulated significantly more biomass on soils conditioned by *G. max* and *M. sativa* compared to the overall mean of all precrops (*p* < 0.001), indicating a positive plant-soil feedback. The growth suppression by *T. aestivum, P. tanacetifolia,* and *B. napus* soils was less pronounced in wild-type plants compared to *bx1* mutant plants. A similar pattern was observed for plant height (**Fig. S2b**). Analysing soil benzoxazinoid concentrations and their degradation products in the soil after harvest confirmed that wild-type plants released significantly more benzoxazinoids than *bx1* mutant plants (**Fig. 1b**). Thus, benzoxazinoid production in maize can convey resistance against negative plant-soil feedbacks.

**Fig. 1.**
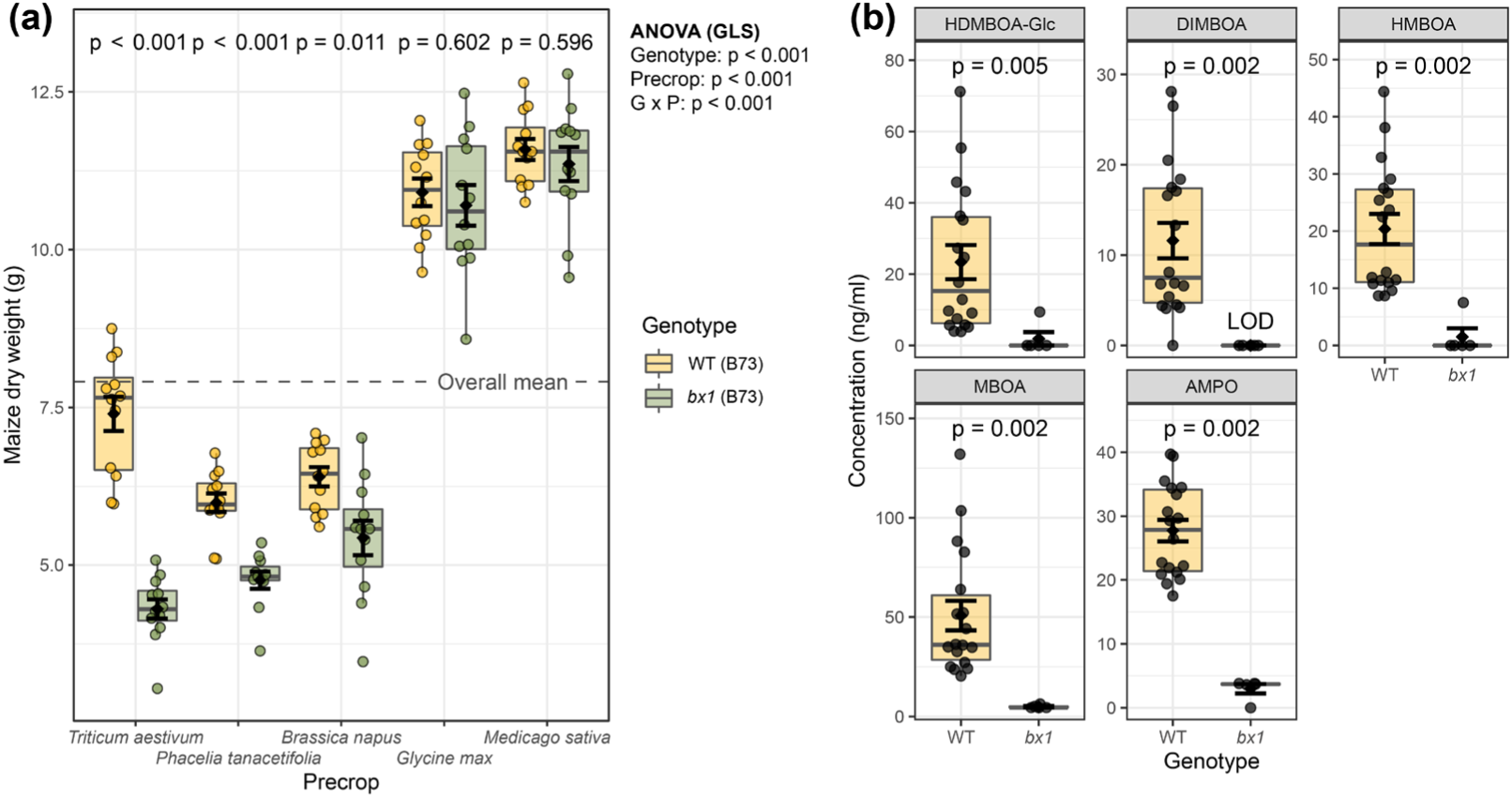
Benzoxazinoid production is associated with resistance to negative plant-soil feedbacks. **(a)** Dry weight of wild-type (WT) or benzoxazinoid-deficient *bx1* mutant maize grown in soils conditioned by five precrop species. Means ± SE, boxplots, and individual datapoints are shown (n = 11-12). ANOVA table and pairwise comparisons of estimated marginal means within each precrop (FDR-corrected *p* values) are provided. The overall mean of all precrops is indicated by the dashed line. **(b)** Soil benzoxazinoid concentration after maize growth of wild-type or benzoxazinoid-deficient *bx1* mutant plants indicated in ng per mL of soil. Means ± SE, boxplots, and individual datapoints are shown (WT: n = 18, *bx1*: n = 5). LOD: below limit of detection. GLS: generalized least squares (linear model). ‘G x P’: interaction between genotype and precrop.

### Benzoxazinoid-dependent resistance to negative plant-soil feedbacks vary across time, soil type, and experiments

Plant-soil feedbacks can be highly context and genotype dependent (Smith-Ramesh & Reynolds, 2017). We thus conducted an additional experiment with a *bx1* mutant in a different genetic background (W22). We also grew wild-type and mutant plants in two different soils (Changins and Q-Matte). We also included B73 plants grown in Changins soil as a positive control. Two plant species with different feedback effects, *T. aestivum* and *M. sativa,* were used to condition the soils (**Fig. S3a**). Three weeks after sowing, the height of wild-type plants was increased compared to *bx1* mutants on *T. aestivum* conditioned Changins soil in both the B73 and the W22 genetic background. Maize plants grew similarly on *M. sativa* conditioned Changins soil, thus confirming that benzoxazinoids increase resistance to negative plant-soil feedbacks (**Fig. 2a**). In contrast to the Changins soil, no difference between wild type and *bx1* mutants in the W22 background was found in Q-Matte soil, illustrating that the suppressive effect induced by *T. aestivum* depends on the soil type (**Fig. 2a**).

**Fig. 2.**
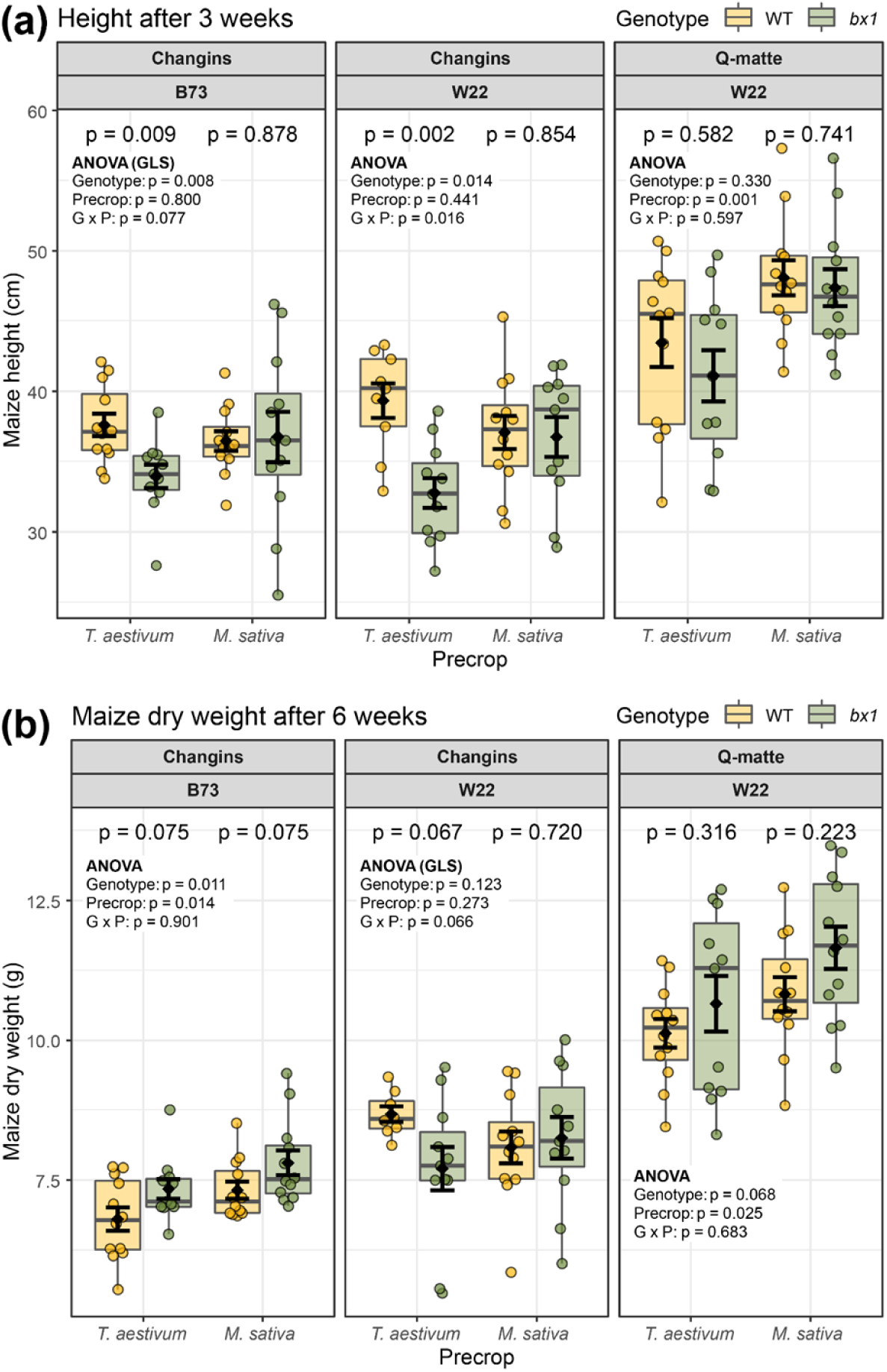
Benzoxazinoid-dependent resistance to negative plant-soil feedbacks is soil-specific and transient. **(a)** Height after three weeks of growth and **(b)** dry weight at harvest of wild-type or benzoxazinoid-deficient *bx1* mutant plants of the maize lines B73 or W22 growing in different soils (Changins, Q-matte) that were previously conditioned by *Triticum aestivum* or *Medicago sativa*. Means ± SE, boxplots, and individual datapoints are shown (n = 8-12). ANOVA table and pairwise comparisons of estimated marginal means within each precrop (FDR-corrected *p* values) are provided. GLS: generalized least squares (linear model). ‘G x P’: interaction between genotype and precrop.

Six weeks after sowing, the differences in height between wild-type and mutant plants were less pronounced, and even reversed in the B73 background in the Changins soil (**Fig. S3b**). Dry weight patterns for W22 were as expected from the early height data, with the *bx1* mutant accumulating less biomass than the wild type in *T. aestivum* conditioned Changins soil, and no difference in *M. sativa* conditioned Changins soil as well as Q-Matte soil (**Fig. 2b**). No clear conditioning effects on biomass were observed in the B73 background. Thus, while benzoxazinoids increase resistance to negative plant-soil feedbacks, the strength of the effects varies across time, different soil types and experiments.

To better capture the variation in feedback resistance, we conducted further experiments with *T. aestivum* conditioned soil from Changins (**Fig. S1, Fig. S4**). First, we performed a detailed time course analysis to better understand the temporal variation in feedback resistance. No differences in germination were observed between genotypes (**Fig. 3a**). After 18 days of growth, wild-type plants grew significantly taller than *bx1* mutant plants in *T. aestivum* conditioned soil. The effect was most pronounced 27 days after sowing. Chlorophyll contents and dry weight were increased in wild-type compared to *bx1* mutant plants at day 27 (**Fig. 3b/c**). Thus, feedback effects appear two weeks after sowing maize and are clearly visible four weeks after sowing. Based on these results, we set the feedback phase to four weeks in all further experiments.

**Fig. 3.**
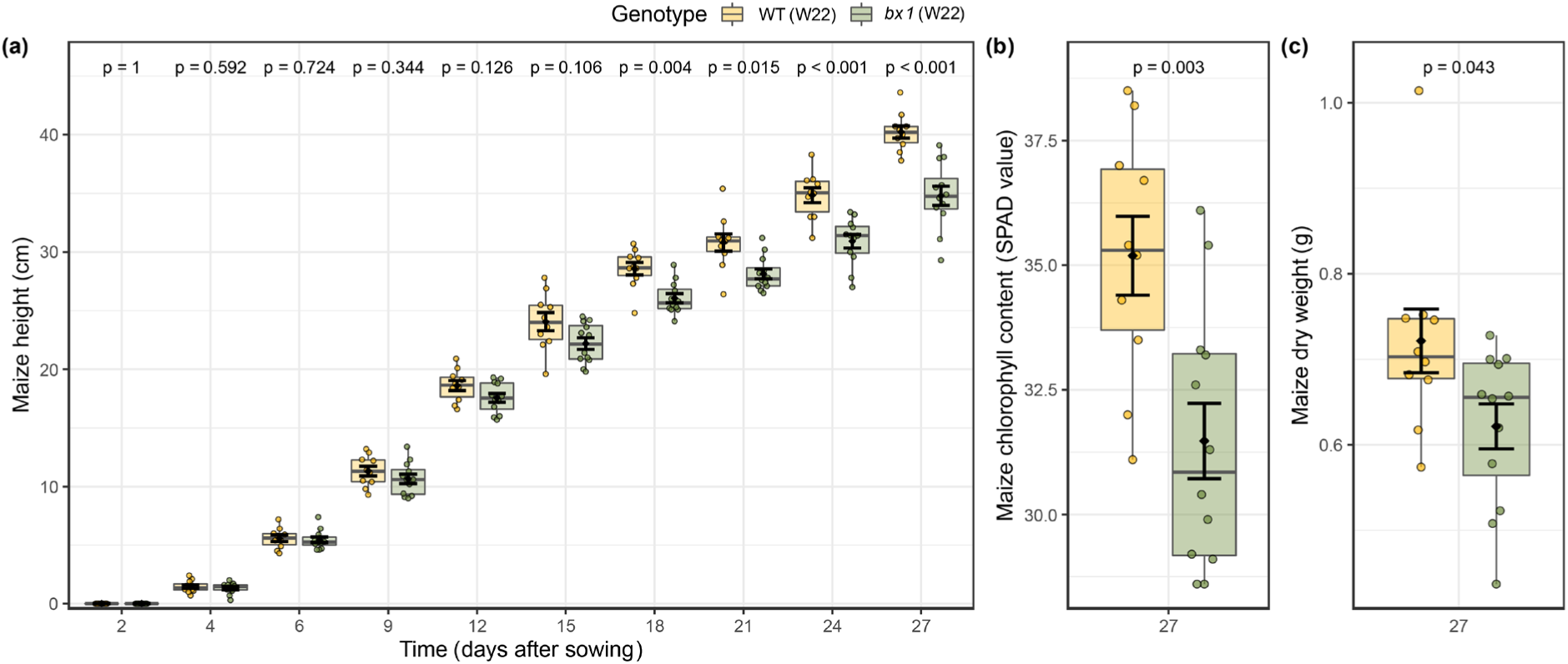
Benzoxazinoid-dependent resistance to negative plant-soil feedbacks appears in early seedling growth. **(a)** Time series of plant height, **(b)** end point chlorophyll content, and **(c)** dry weight at harvest of wild-type or benzoxazinoid-deficient *bx1* mutant plants grown in soils that were conditioned by *Triticum aestivum*. Means ± SE, boxplots, and individual datapoints are shown (n = 10-12). Statistical significance is indicated as *p* values computed by Welch’s two-sample *t*-test. *p* values were adjusted for multiple testing (FDR) in **(a).**

### Benzoxazinoids in the soil increase resistance to negative plant-soil feedbacks

To test if benzoxazinoids act in the soil, we complemented the soil of *bx1* mutant plants with a benzoxazinoid mixture typical for young maize seedlings (**Fig. S5a/b**). We then compared the performance of wild-type and *bx1* mutant plants grown in *T. aestivum* conditioned soil with and without benzoxazinoid supplementation. Three different complementation concentrations were applied in three consecutive experiments.

There was considerable variation in the observed phenotypic effects across experiments. In each experiment, at least one out of three measured plant performance parameters was enhanced in wild-type plants compared to *bx1* mutant plants growing in *T. aestivum* conditioned soil (**Fig. 4**). In each case, benzoxazinoid supplementation rescued the wild-type phenotypes, either fully or partially. The clearest effect was observed for chlorophyll contents (**Fig. 4**). Interestingly, even though the applied benzoxazinoid concentrations varied by two orders of magnitude, we observed complementation effects in all three experiments (**Fig. 4**). Taken together these results suggest that despite considerable variation, resistance to negative plant-soil feedbacks can be explained by benzoxazinoid release into the soil.

**Fig. 4.**
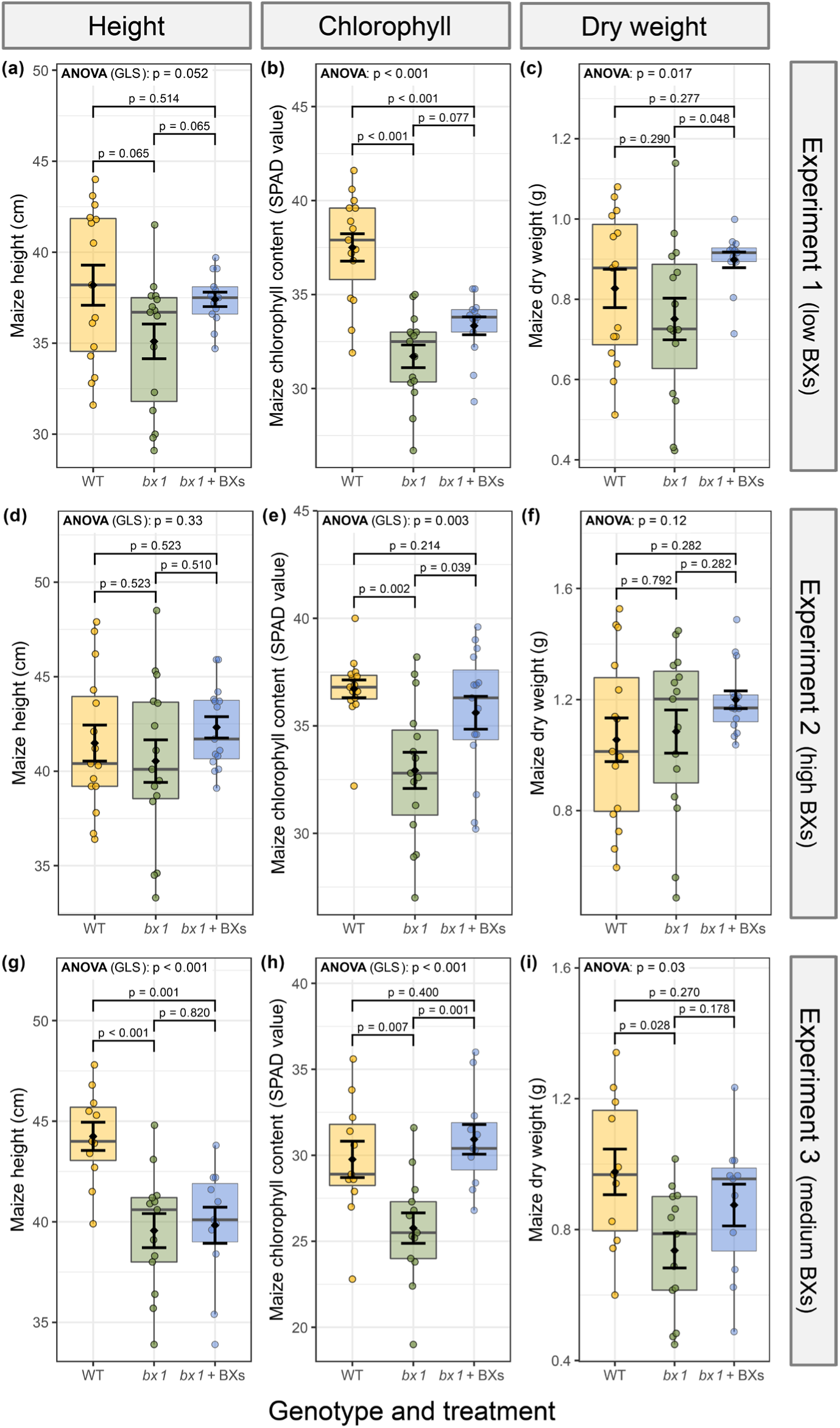
Benzoxazinoid-dependent resistance to negative plant-soil feedbacks is partially associated with benzoxazinoids in the rhizosphere. For all three replications of this experiment height, chlorophyll content, and dry weight of wild-type plants, benzoxazinoid-deficient *bx1* mutant plants, or *bx1* plants complemented with benzoxazinoids grown in soils that were previously conditioned by *Triticum aestivum*. Means ± SE, boxplots, and individual datapoints are shown. ANOVA table and pairwise comparisons of estimated marginal means between all three treatments (FDR-corrected *p* values) are provided. Experiment 1, 2 and 3 were complemented with low, high, and medium amounts of benzoxazinoids. Experiment 1: n = 13-15, experiment 2: n = 15, experiment 3: n= 11-13. GLS: generalized least squares (linear model). ‘G x P’: interaction between genotype and precrop. BXs: benzoxazinoids.

### Soil microbiota drive benzoxazinoid-dependent resistance to negative plant-soil feedbacks

To investigate the role of soil microbiota in benzoxazinoid-mediated resistance against negative plant-soil feedbacks, we X-ray sterilized part of the conditioned soil and grew wild-type and *bx1* mutant plants in the soils. A microbial re-inoculation treatment was included to control for changes in soil chemical and physical properties that may result from the sterilization treatment (Berns *et al*., 2008). As expected, wild-type plants outperformed *bx1* mutants across experiments in one or several performance parameters when growing in *T. aestivum* conditioned soil (**Fig. 5**). All resistance effects were lost in sterilized soil. Re-inoculation restored all resistance effects in experiment 2. In experiments 1 and 3, only tendencies for restored resistance effects were found in re-inoculated soil. Thus, benzoxazinoid-mediated resistance is mediated by elements that are labile to sterilization. At least in some cases, soil biota can account for these effects, as reinoculation with a small quantity of soil is sufficient to restore benzoxazinoid-mediated resistance.

**Fig. 5.**
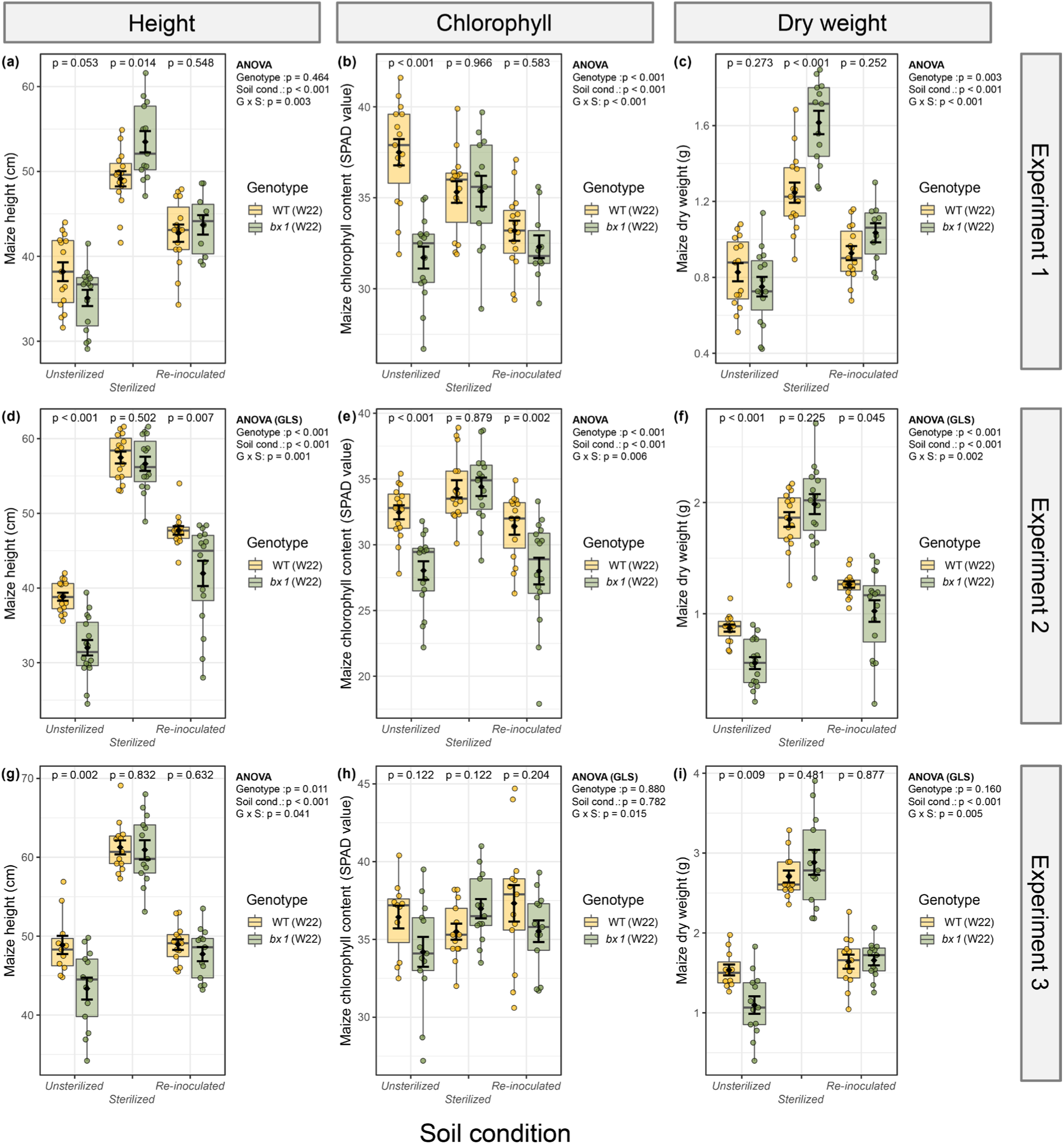
Benzoxazinoid-dependent resistance to negative plant-soil feedbacks can depend on soil biota. For all three replications of this experiment height, chlorophyll content, and dry weight of wild-type or benzoxazinoid-deficient *bx1* mutant plants grown in *Triticum aestivum* conditioned soil that was either unsterilized, sterilized, or sterilized and re-inoculated with unsterilized soil. Means ± SE, boxplots, and individual datapoints are shown. ANOVA table and pairwise comparisons of estimated marginal means between all three treatments (FDR-corrected *p* values) are provided. Experiment 1: n = 10-15, Experiment 2: n = 15-16, Experiment 3: n= 11-13. GLS: generalized least squares (linear model). ‘G x S’: interaction between genotype and soil condition.

To further examine the role of soil biota in benzoxazinoid-dependent plant-soil feedbacks, we conducted an additional inoculation experiment. We sterilized *M. sativa* conditioned soil and inoculated it with either *M. sativa* or *T. aestivum* soil. In unsterilized *M. sativa* conditioned soil, wild-type and *bx1* mutant plants grew similarly well, as observed before (**Fig. 6**). In sterilized soils, the *bx1* mutant outperformed wild-type plant growth. This effect disappeared when the soil was inoculated with *M. sativa* soil (**Fig. 6**). When the soil was inoculated with *T. aestivum* biota, wild-type plants outperformed *bx1* mutant plants. This reciprocal transplant experiment shows that the negative effects of *T. aestivum* soil biota can be overcome by benzoxazinoids.

**Fig. 6.**
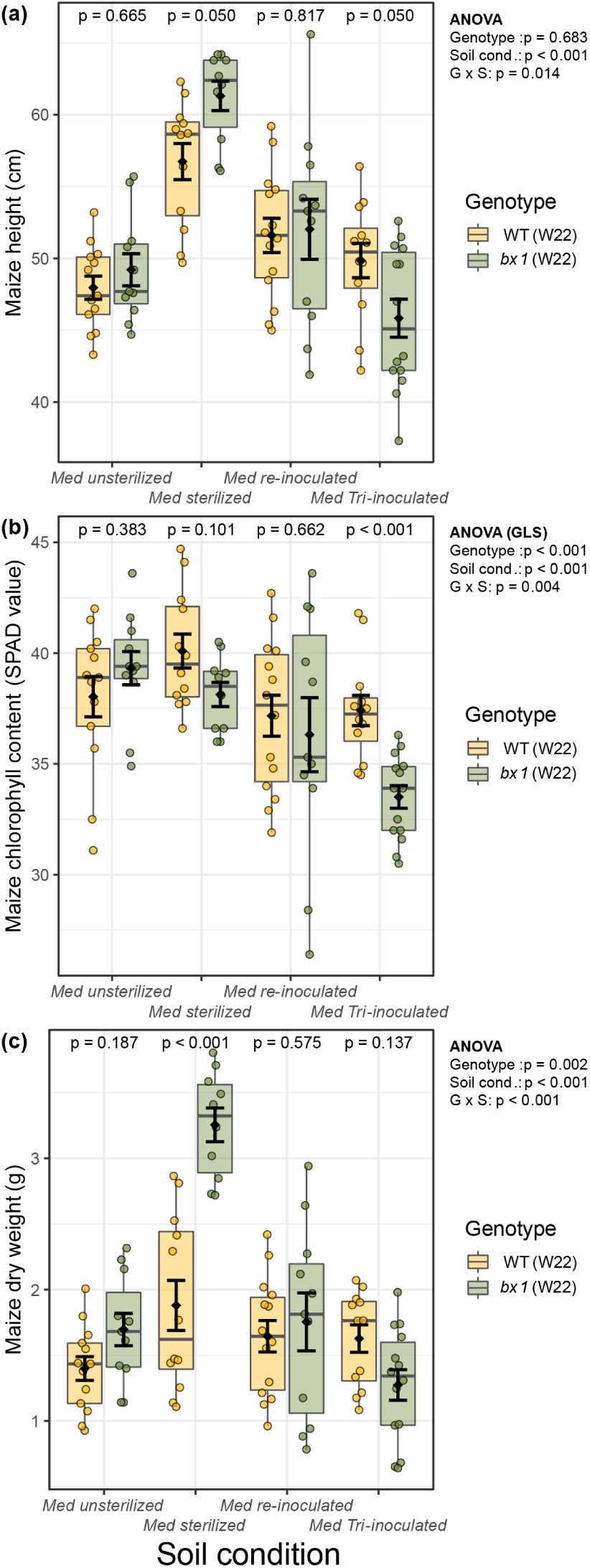
Benzoxazinoid-dependent resistance to negative plant-soil feedbacks depends on precrop-specific soil biota. **(a)** Height, **(b)** chlorophyll content, and **(c)** dry weight of wild-type or benzoxazinoid-deficient *bx1* mutant plants grown in *Medicago sativa* conditioned soil that was either unsterilized, sterilized, sterilized and re-inoculated with unsterilized *Medicago sativa* soil (Med-inoculated), or sterilized and re-inoculated with unsterilized *Triticum aestivum* soil (Tri-inoculated). Means ± SE, boxplots, and individual datapoints are shown (n = 10-14). ANOVA table and pairwise comparisons of estimated marginal means between all three treatments (FDR-corrected *p* values) are provided. GLS: generalized least squares (linear model). ‘G x S’: interaction between genotype and soil condition.

## Discussion

Plant-soil feedbacks have a major impact on plant performance. How plants resist negative plant-soil feedbacks is not well known. In this study we demonstrate that benzoxazinoids can help plants to cope with variable negative plant-soil feedbacks. This effect, is at least partially, mediated by the interaction between benzoxazinoids and soil microbiota. Below we discuss the underlying mechanisms and agroecological implications of our findings.

Plant-soil feedbacks can be triggered through exuded secondary metabolites and their capacity to change root-associated microorganisms (Hu *et al*., 2018b; Yu *et al*., 2021). To what extent root secondary metabolites can protect plants from negative plant-soil feedbacks is unknown. Our results demonstrate that benzoxazinoid exudation into the rhizosphere can mitigate negative plant-soil feedbacks. This effect was found in two maize lines and was most pronounced for early performance; an important trait in crop cultivation (Ellis, 1992; Steege *et al*., 2005; Shi *et al*., 2020). Benzoxazinoids are known to shape the root and rhizosphere microbiome and suppress particular soil pathogens (Wilkes *et al*., 1999; Martyniuk *et al*., 2006; Cadot *et al*., 2021b), therefore benzoxazinoid-mediated resistance could be driven by the mitigation of adverse impacts of soil born plant pathogens, which are known to be capable of massively reducing seedling performance (Packer & Clay, 2000). Indeed, sterilization resulted in the disappearance of the negative effect and the capacity of benzoxazinoids to improve plant performance, and (re)-inoculation with soil biota partially re-established the effects. Given the wide range of metabolites plants can employ to modulate root microbiota and establish their own, often beneficial microbial communities (Pang *et al*., 2021), we propose that this form of soil conditioning may be a widespread mechanism that protects plants from growth suppression by other plants. To what extent such conditioning may be costly by reversing positive feedback effects remains to be established.

Plant-soil feedback effects are known to be highly context dependent, rendering them variable to a point where seemingly stochastic patterns are observed. Plant-soil feedbacks are known to depend on the growth environment (Schittko *et al*., 2016), soil origin, above-ground herbivores, soil microbes, as well as temperature and soil moisture (Long *et al*., 2019; Cadot *et al*., 2021a). Small variations in abiotic and biotic parameters may have contributed to the variation within and between experiments that we observed in our study, even under controlled conditions (Wei *et al*., 2019). Despite this variation, we observed a remarkable consistency in the directionality of our effects, suggesting that, while quantitatively variable, the net protective effect of benzoxazinoids towards negative plant-soil feedbacks is relevant for plant performance. Nevertheless, the benefits of benzoxazinoid exudation is likely to depend on the soil environment (Cadot *et al*., 2021a). While negative plant-soil feedbacks are observed in one soil, they are absent in another soil, and thus, no protection is afforded by benzoxazinoids in this second situation. Interestingly, this second soil, Q-matte, has previously been shown to be incapable of provoking benzoxazinoid dependent plant-soil feedbacks on successor plants (Cadot *et al*., 2021a). Experiments with additional soils will show how important benzoxazinoid-mediated resistance is in an agroecological context.

Crop rotations have been incorporated into agricultural practices for centuries to lower negative effects of crops, such as accumulation of species-specific soil-borne pathogens or nutrient depletion (van der Putten *et al*., 2013). Only recently, cultivar-specific feedbacks within agricultural plant-soil feedbacks have been demonstrated (Wagg *et al*., 2015; Carrillo *et al*., 2019; Cadot *et al*., 2021a; Awodele & Bennett, 2022). The mechanisms responsible for tolerating a given precrop is largely unexplored. In our work, we find that one single group of secondary metabolites controlled by one single gene (*Bx1*) determines the resistance of maize against negative plant-soil feedbacks. Given that the same metabolites can increase agricultural productivity of the following crop (Gfeller *et al*., 2022), this makes the genes involved in biosynthesis and exudation of such metabolites a potential breeding target for superior crop rotations. Many maize lines already produce substantial amounts of benzoxazinoids in their roots, but substantial genetic variation is commonly observed (Handrick *et al*., 2016). It should thus be possible to develop cultivars that are particularly suited to crop rotations or that may deliver better performance following specific preceding crops. Broader field experiments will be needed to quantify the potential of optimized benzoxazinoid release to promote sustainable crop production by improving yields and food quality while reducing inputs.

## Conclusion

Plants strongly interact with the soil, where the release of secondary metabolites has a strong effect on soil biota (Sasse *et al*., 2018). Our study shows that such exudation may increase crop rotation stability by reducing negative plant-soil feedbacks. The use of agroecological plant-soil feedbacks has been proposed as a possible way towards more sustainable systems (Mariotte *et al*., 2018) and with our work we provide an additional mechanism to apply this concept. As the release of diverse secondary metabolites into the rhizosphere is a common plant trait (Baetz & Martinoia, 2014), studying their effect on crop rotations offers a big reservoir of possible mechanisms to make agriculture more sustainable through plant-soil feedbacks (Mariotte *et al*., 2018).

## Acknowledgement

We thank Florian Enz, Sophie Gulliver, and Pascal Wyss for their assistance with the growth and maintenance of plants. Further, we are grateful to Pierre Mateo for helpful advice on benzoxazinoid purification and analytics. This work was supported by the Swiss National Science Foundation (Grant. Nr. 192564) and the Interfaculty Research Collaboration “One Health” of the University of Bern.

## Author contributions

V.G. and M.E. designed research; V.G. and L.T. performed research; V.G. and M.E. analysed data; and V.G. and M.E. wrote the first draft of the paper. All authors revised the paper.

## Data availability

All data generated for this study can be downloaded from the DRYAD repository (doi to be inserted). R codes to reproduce statistical analysis and visualizations is available under (https://github.com/ValentinGfeller/Precrop_paper).

## Supplementary Figures

**Fig. S1.**
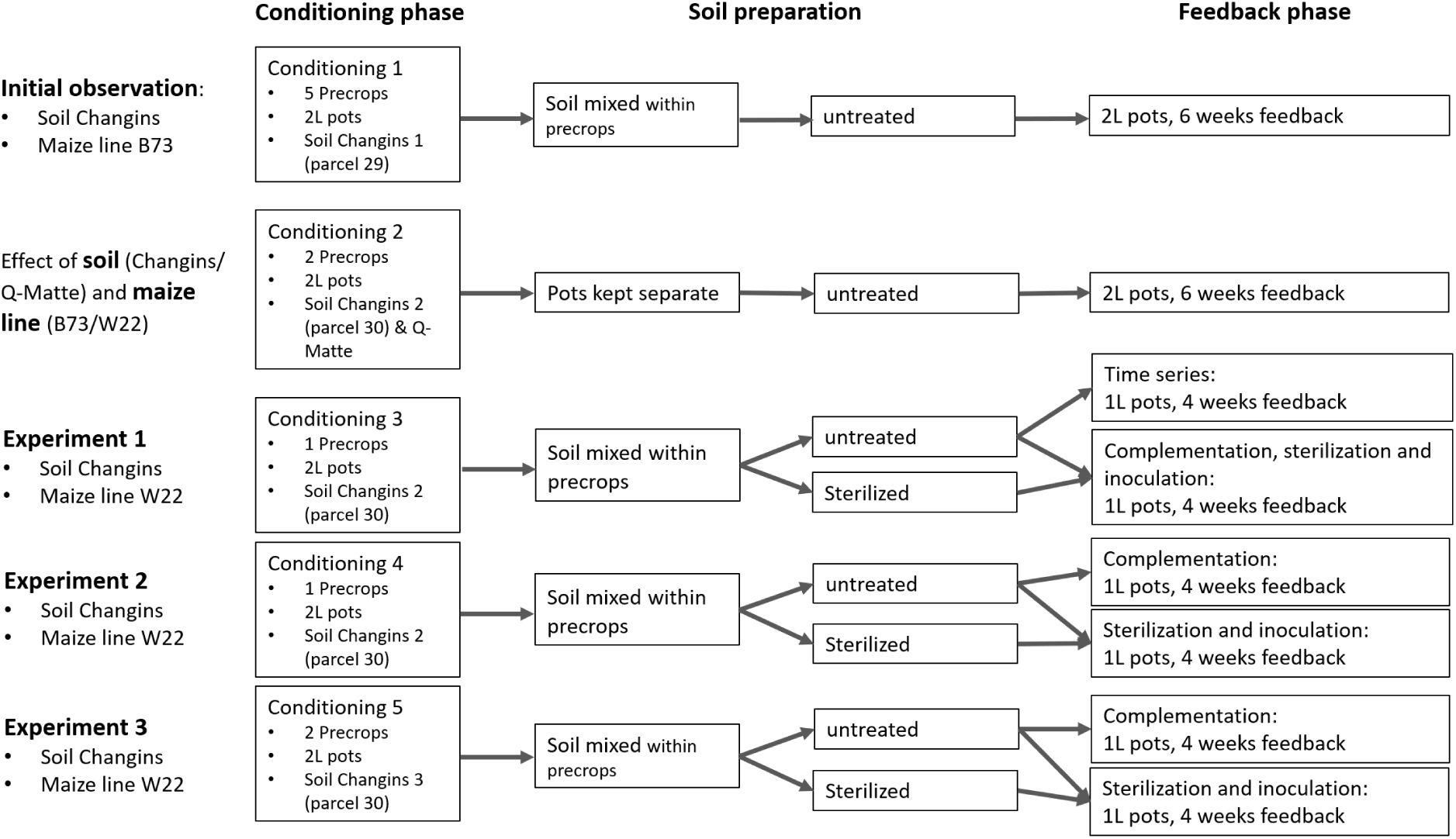
Experimental setup. All experiments are listed and specification concerning the conditioning and the feedback phase are indicated. Further, soil preparation between conditioning and feedback is shown.

**Fig. S2.**
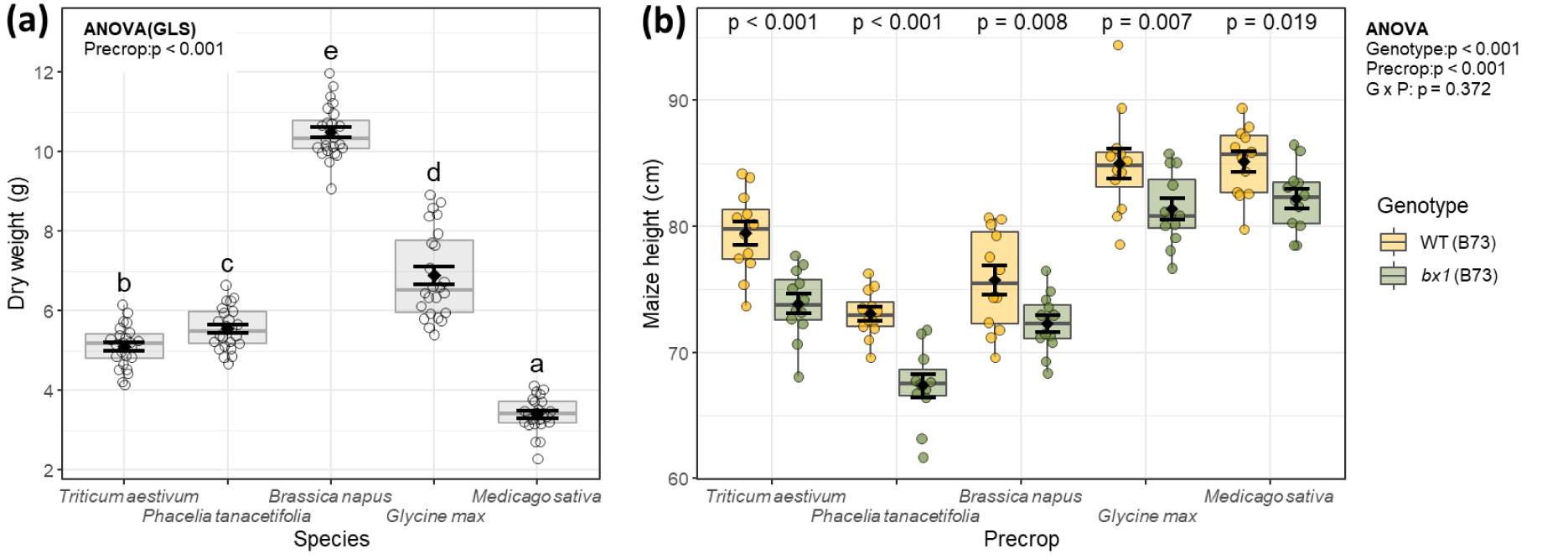
Precrop weight and maize height of initial precrop screening experiment. **(a)** Dry weight of precrops grown to condition the soil for the initial experiment and **(b)** height of wild-type or benzoxazinoid-deficient *bx1* mutant maize grown in soils conditioned by five precrop species. **(a)** Means ± SE, boxplots, and individual datapoints are shown (n = 24). ANOVA table and compact letter display of all pair-wise comparisons (Significance-level: FDR-corrected *p* < 0.05) of estimated marginal means are provided. **(b)** Means ± SE, boxplots, and individual datapoints are shown (n = 11-12). ANOVA table and pairwise comparisons of estimated marginal means within each precrop (FDR-corrected *p* values) are provided. ‘G x P’: interaction between genotype and precrop.

**Fig. S3.**
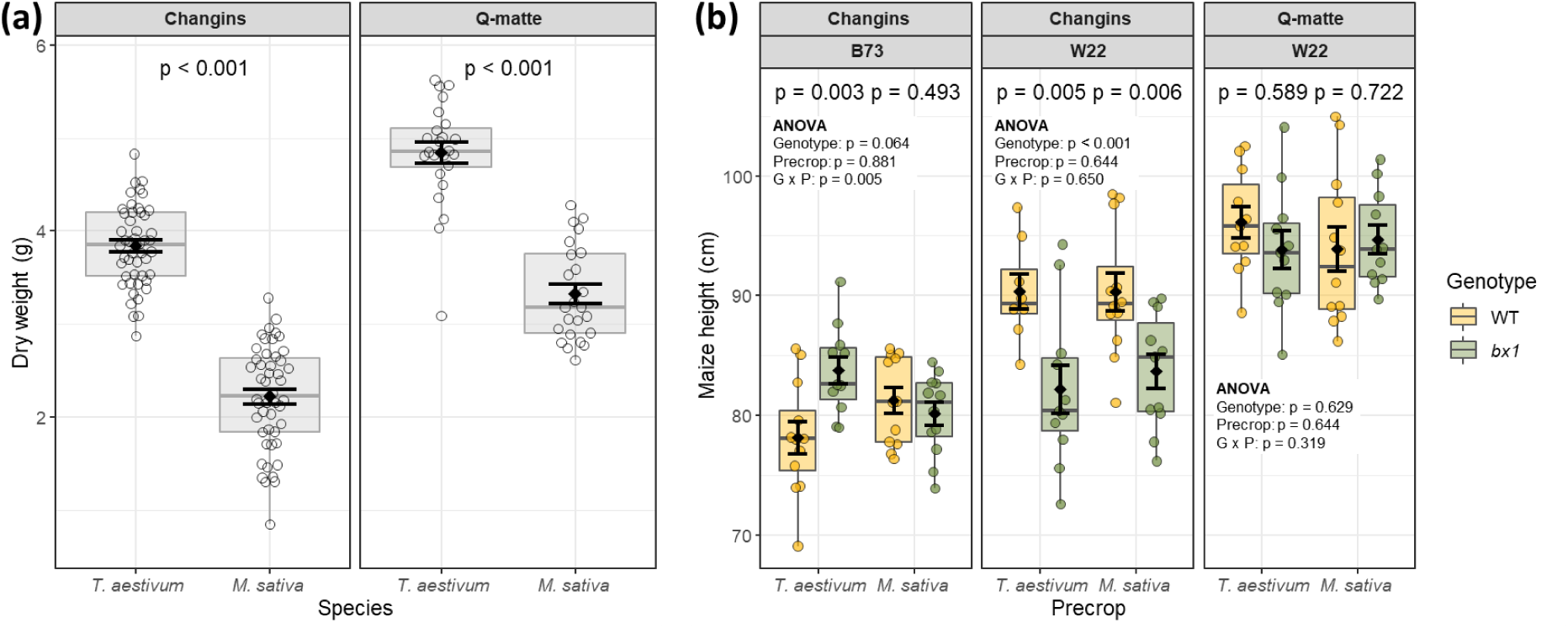
Precrop weight and maize height of experiment comparing different soil origins and maize lines. **(a)** Dry weight of precrop species grown in two different soils (Changins and Q-matte) and **(b)** height at harvest of wild-type or benzoxazinoid-deficient *bx1* mutant plants of the maize lines B73 or W22 growing in different soils that were conditioned by *Triticum aestivum* or *Medicago sativa*. **(a)** Means ± SE, boxplots, and individual datapoints are shown (Changings: n = 48, Q-matte: n= 24). Statistical significance is indicated as *p* values computed by Welch’s two-sample *t*-test. **(b)** Means ± SE, boxplots, and individual datapoints are shown (n = 8-12). ANOVA table and pairwise comparisons of estimated marginal means within each precrop (FDR-corrected *p* values) are provided. GLS: generalized least squares (linear model). ‘G x P’: interaction between genotype and precrop.

**Fig. S4.**
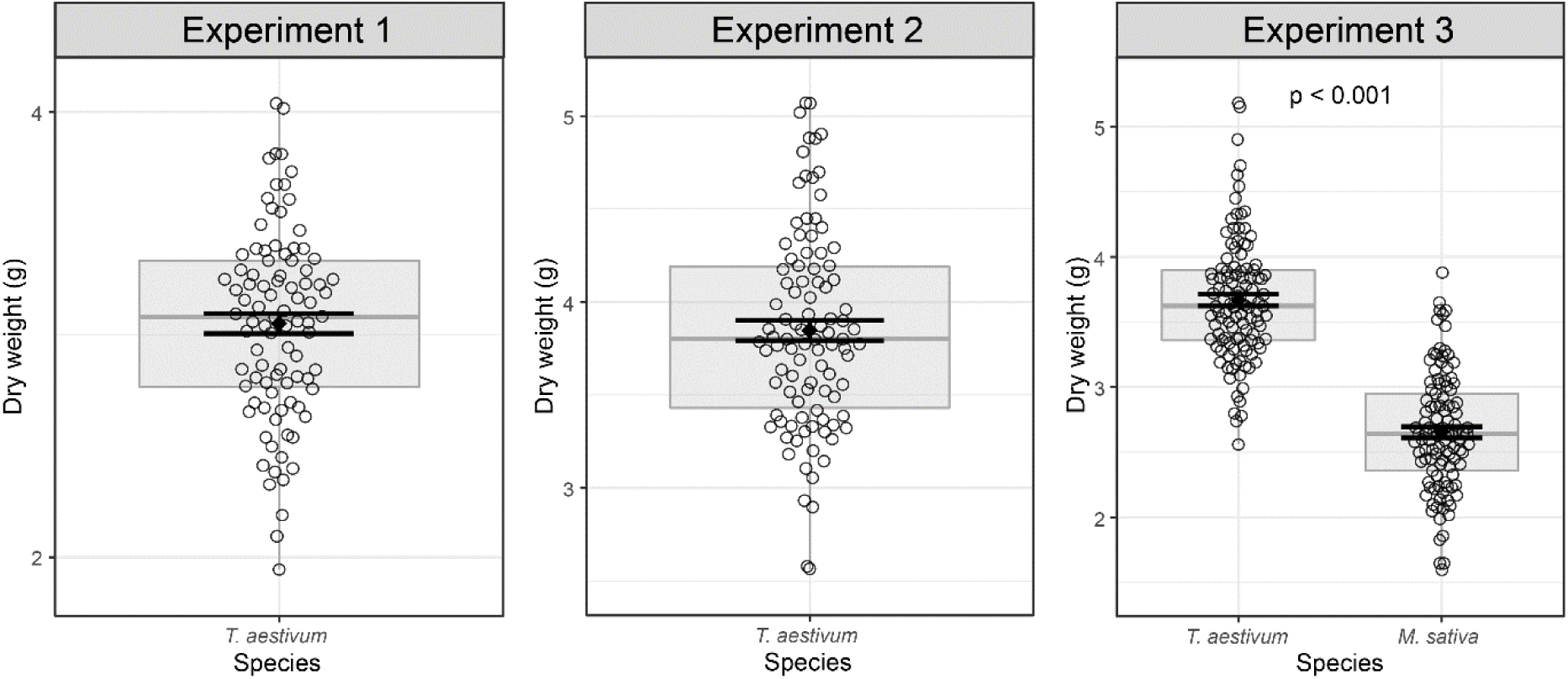
Dry weight of precrop species grown to condition the soil for replicated experiments 1-3. Means ± SE, boxplots, and individual datapoints are shown. For experiment 3 statistical significance is indicated as *p* values computed by Welch’s two-sample *t*-test. Experiment 1: *T. aestivum* n = 92; experiment 2: *T. aestivum* n = 98; experiment 3: *T. aestivum* n = 110, *M. sativa* = 109.

**Fig. S5.**
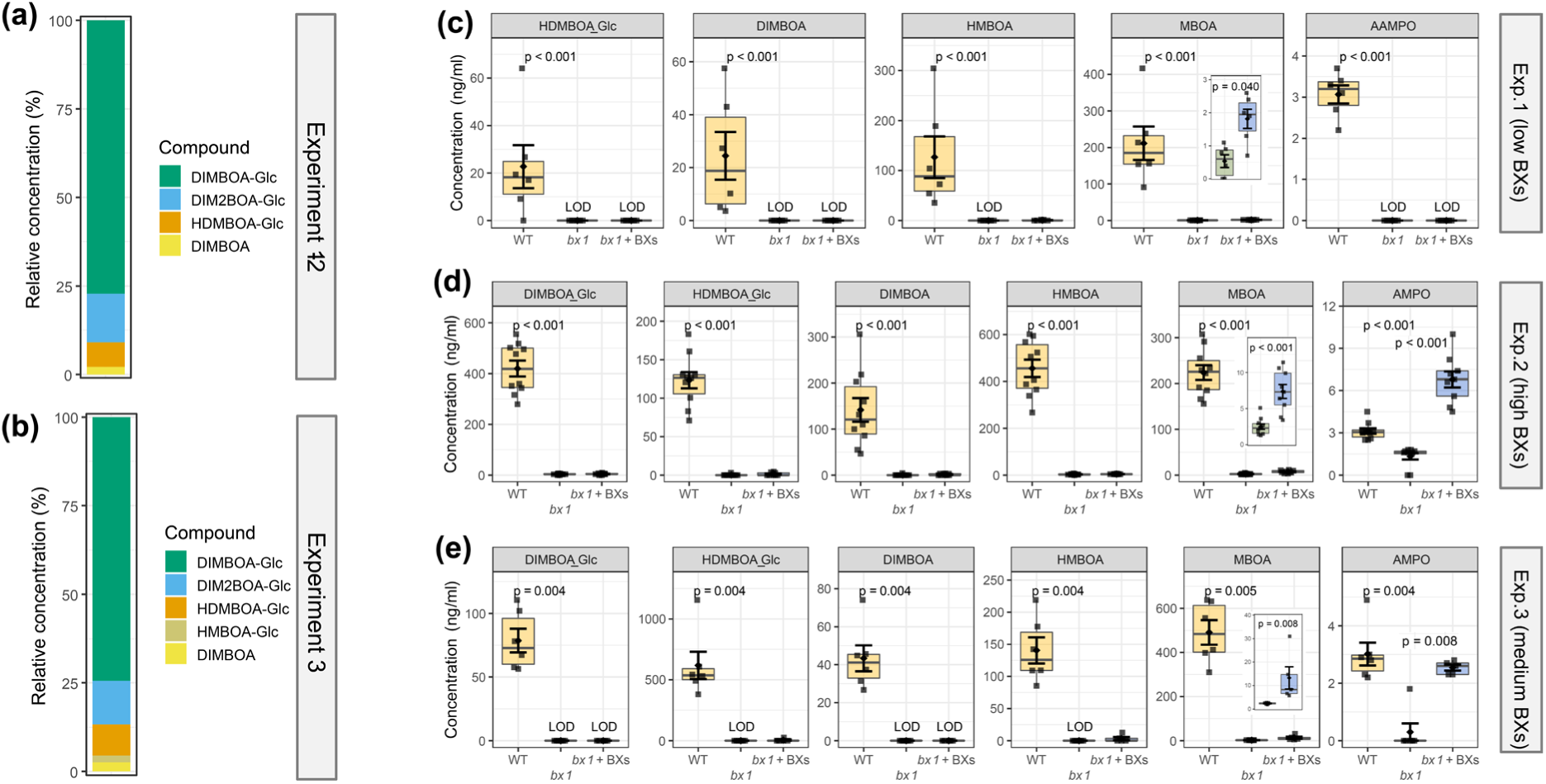
Characterization of benzoxazinoid complementation. **(a, b)** relative abundance of single benzoxazinoids in mixture purified from four days germinated maize kernels and applied for complementation. **(c-e)** Soil benzoxazinoid concentration after maize growth of wild-type, benzoxazinoid-deficient *bx1* mutant plants, or *bx1* plants complemented with benzoxazinoids indicated in ng per mL of soil. Means ± SE, boxplots, and individual datapoints are shown. Provided *p* values for the comparison between wild-type and *bx1* plants were computed by Wilcoxon rank-sum tests and corrected for multiple testing (FDR). If significant, p values for the comparison between *bx1* plants and complement *bx1* pants are also provided. Experiment 1, 2 and 3 were complemented with low, high, and medium amounts of benzoxazinoids. Experiment 1: n = 6, experiment 2: n = 9-10, experiment 3: n= 5-6. LOD: below limit of detection. BXs: benzoxazinoids.

